# PSI Atlas: a comprehensive knowledgebase of plant self-incompatibility

**DOI:** 10.1101/2025.01.07.631397

**Authors:** Chen Wang, Hong Zhao, Hongkui Zhang, Sijie Sun, YongBiao Xue

## Abstract

Self-incompatibility (SI) is an important genetic mechanism in angiosperms that prevents inbreeding and promotes outcrossing. Although significant progress has been made in understanding SI, its molecular underpinnings and evolutionary origins remain elusive in many plant families. In eudicots, SI is typically regulated by a single *S*-locus with diverse *S*-haplotypes, while in grasses, it is controlled by two separate *S* and *Z* loci. Genome assemblies of SI plants have greatly facilitated the identification and evolutionary analysis of *S*-loci. However, a comprehensive database that integrates information on the diverse *S*-loci has been lacking. To fill this gap, we create the Plant Self-Incompatibility Atlas (PSIA, http://www.plantsi.cn), a knowledgebase that offers an extensive compilation of plant SI, including genomic resources for assembled SI species, the origins and evolution of *S* genes, and the molecular mechanisms of the eight known SI types. In our most recent release, we have obtained more than 500 genome assemblies across 469 SI species. We have also collected 1275 nucleotide and 1130 protein sequence accessions of *S* genes from public databases, with a total of 3095 *S* genes manually identified and curated. PSIA not only thoroughly explores the *S*-locus information of the assembled SI species but also enables users to efficiently browse, perform BLAST searches, analyze, and download *S* genes. Additionally, PSIA acts as a comprehensive platform for comparative genomic studies of *S*-loci, aiding in the exploration of the dynamic processes involved in the origin, loss, and regain of SI. Consequently, PSIA is poised to significantly enhance our understanding of angiosperm SI and offer new perspectives on plant mating systems.

## Introduction

Self-incompatibility (SI), found in over 40% of flowering plants [1], is a reproductive mechanism that prevents a fertile seed plant from producing a zygote through self-fertilization. As a strict intraspecific barrier, SI is crucial for preventing inbreeding and promoting outcrossing, which are essential for maintaining genetic diversity and the adaptive survival of populations [2]. In eudicots, SI is typically controlled by a highly polymorphic multi-allelic *S*-locus [3], while in grasses, it is governed by two independently inherited, polymorphic *S* and *Z* loci [4]. These loci are characterized by the tight linkage of distinct gene types that control the *S* or *Z* specificities in the pistil and pollen. In eudicot SI, self-pollen incompatibility (SPI), leading to self-pollen rejection, is triggered by the self-recognition of pistil and pollen *S* determinants encoded by the same *S* haplotype. In contrast, in grasses, SPI occurs only when both the *S* and *Z* haplotypes match. SI systems can be categorized into homomorphic (gametophytic and sporophytic) and heteromorphic (heterostyly) types based on their association with floral morphology [2, 3, 5-7]. Gametophytic SI (GSI), where self-pollen rejection is determined by its own genotype, is found in 17-25 families, while sporophytic SI (SSI), where self-pollen rejection is determined by its parental genotype, is found in a few families, including Brassicaceae, Asteraceae, Convolvulaceae, and Betulaceae [8-12]. Heterostyly, which mainly exists in 28 families such as Primulaceae, Turneraceae, Linaceae, and Oleaceae [13-16], combines morphological and physiological incompatibility to prevent within-morph fertilization.

Given the significant variation in the molecular mechanisms underlying SI systems across different families, they can be further divided into eight types. These include gametophytic type-1 SI, controlled by the pistil *S S-RNase* and pollen *S SLF*, found in Plantaginaceae [17-19], Solanaceae [20-22], Rosaceae [23-25], and Rutaceae [26]; sporophytic type-2 of Brassicaceae, governed by *SRK* and *SP11*/*SCR* [8, 9]; gametophytic type-3 of *Papaver rhoeas*, determined by *PrsS* and *PrpS* [27, 28]; heterostyly type-4 of Primulaceae, defined by a hemizygous *S*-locus encoding *CYP, GLO2, KFB, CCM*, and *PUM* [13]; heterostyly type-5 of Turneraceae, controlled by *TsSPH1, TsYUC6*, and *TsBAHD* [14]; gametophytic type-6 of Poaceae, by *HPS10-S/Z* and *DUF247I/II-S/Z* [4, 29, 30]; heterostyly type-7 of Linaceae, by *LtTSS1* and *LtWDR44* [15]; and type-8 of Oleaceae, by *GA2ox* [16].

Recent progress in high-quality genome assemblies has significantly benefited the identification and evolutionary studies of *S*-loci. However, a comprehensive database integrating the information and knowledge of the diverse *S*-loci is still lacking. PSI Atlas (PSIA) fills this gap as the first extensive knowledgebase on plant SI, meticulously curated from molecular mechanism studies and extensive public genome resources.

## Implementation

PSIA runs on a Linux-based Apache (https://httpd.apache.org) web server using PHP (A popular general-purpose scripting language especially suited to web development; https://www.php.net) as the backend. The web interface is built with Bootstrap5 (A powerful, extensible, and feature-packed frontend toolkit; https://getbootstrap.com), HTML5, CSS, and JavaScript. MySQL (http://www.mysql.org) has been adopted as the relational database management system to store information. All codes are developed using Visual Studio (https://visualstudio.microsoft.com), a powerful and versatile integrated development environment (IDE) for software developers and teams. To provide stable web services, PSIA is hosted on the Elastic Cloud Server (ECS) of HUAWEI CLOUD (https://activity.huaweicloud.com). The website has been tested to ensure its functionality across various operating systems and web browsers such as Google Chrome, Firefox, and Microsoft Edge.

## Database content and usage

### Integration of genome resources

Approximately 40% of flowering plant species from at least 100 families possess SI [1]. Recent studies have identified eight distinct molecular types of SI at the family level, which encompass eleven families containing a variety of crops, fruits, ornamental and model plants (Figure 1A and 1B). For instance, potato, tomato, pepper and tobacco in Solanaceae (type-1), strawberry and apple in Rosaceae (type-1), citrus in Rutaceae (type-1), snapdragon in Plantaginaceae (type-1), cabbage in Brassicaceae (type-2), opium poppy in Papaveraceae (type-3), perennial ryegrass in Poaceae (type-6), heterostyly model plants in Primulaceae (type-4), Turneraceae (type-5), Linaceae (type-7) and Oleaceae (type-8). Although high-quality genome assemblies of these species have been published recently, most of them overlook the annotation and analyses of *S* genes. Here, we collected and reanalyzed the genomic resources of those species possessing one of the eight SI types. Based on the known molecular types of SI, the *S*-locus information of the assembled SI species will be fully explored in the PSIA.

**Figure 1.**
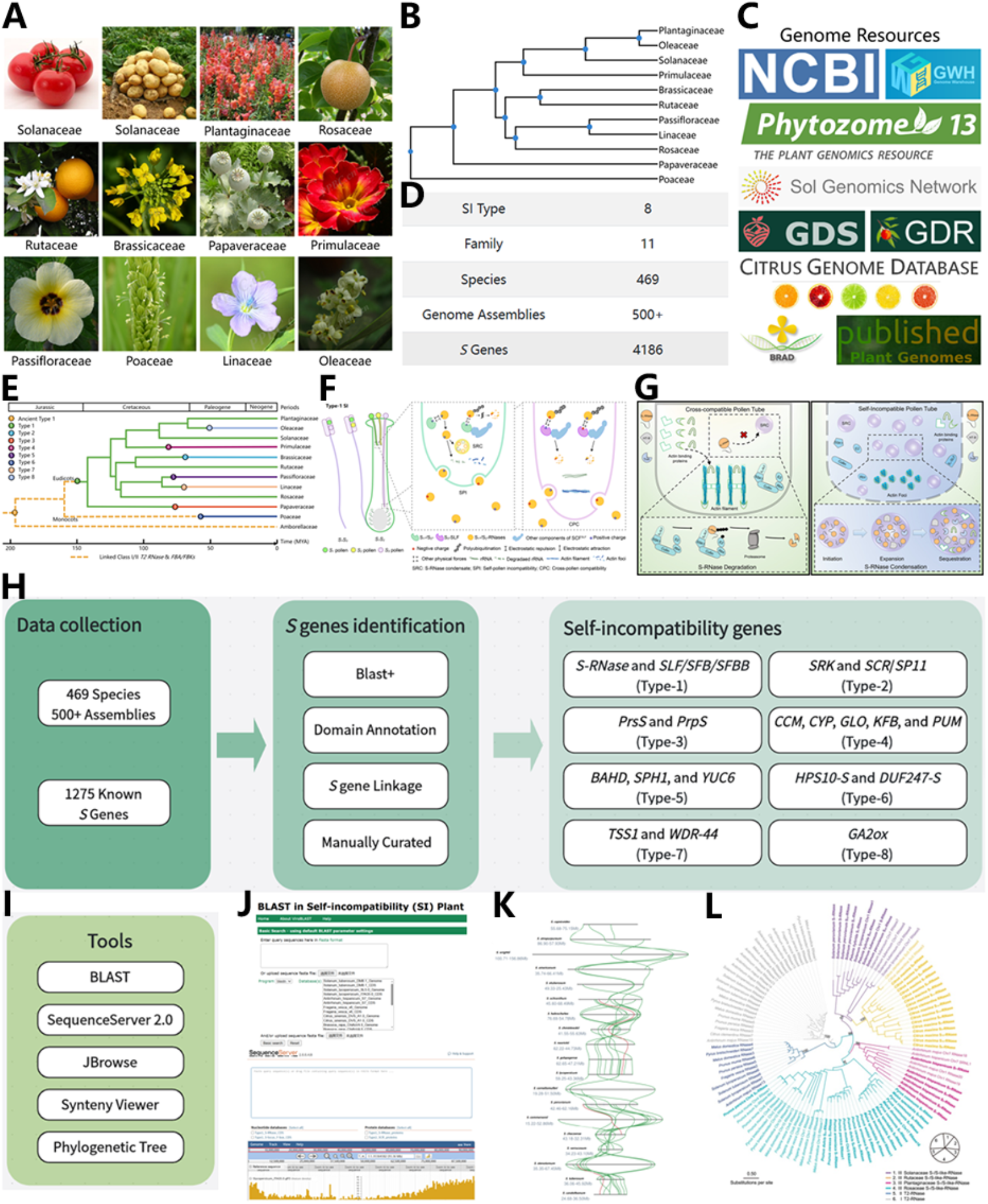
Overview of PSI Atlas. **A**. Representative species of eight SI types. **B**. The phylogenetic relationships of all existing families in PSIA. **C**. Genome resources of SI species. **D**. Overview of SI data in the latest release. **E**. The origin and evolution of the eight SI types. **F**. Diagram illustrating the molecular mechanism of type-1 SI with those of all eight types shown in PSIA. **G**. The reported working model of SI found in the representative species *Petunia hybrida*. **H**. The pipeline of *S* gene identification. **I**. Useful tools of PSIA. **J**. BLAST and JBrowse function of PSIA. **K**. The output of the synteny analysis between the *Solanum S*-loci is shown as an example. **L**. The output of the phylogenetic analysis for S-RNases from the exemplar families such as Plantaginaceae, Solanaceae, Rosaceae and Rutaceae.

PSIA has the most comprehensive genome assembly information of SI plants, with all available genome assemblies of SI species collected from NCBI (National Center for Biotechnology Information, https://www.ncbi.nlm.nih.gov/), NGDC-GWH (Genome Warehouse, https://ngdc.cncb.ac.cn/gwh) [31], Published Plant Genomes (https://www.plabipd.de/plant_genomes_pa.ep), Phytozome v13 (https://phytozome-next.jgi.doe.gov) [32], Sol Genomics Network (https://solgenomics.net) [33], Genome Database for Rosaceae (https://www.rosaceae.org) [34], Genome Database for Strawberry (http://eplant.njau.edu.cn/strawberry/) [35], Citrus Genome Database (https://www.citrusgenomedb.org), Brassicaceae Database (http://www.brassicadb.cn/#/) [36], and other links (such as data availability links in papers, figshare, and google drive) (Figure 1C). In the current release, more than 500 genome assemblies covering 469 SI species were obtained (Figure 1D; Supplemental Figure S1). Besides, our database will be regularly updated to incorporate the latest sequenced genomes of SI plants.

### Knowledge of SI origin, evolution, molecular mechanisms and working models

To facilitate a systematical study of SI, PSIA incorporates fundamental knowledge of plant SI, including its origin and evolution, molecular mechanisms and the reported working models. Several SI systems have been identified in eudicots, with type-1 being the most ancient and widely adopted, while type 2-8 are family-specific. Using a combination of phylogenomic and genetic analyses, Zhao et al., (2022) uncovered a single origin traced back to the most recently common ancestor (MRCA) of eudicots for S-RNase, *SLF/SFB/SFBB* and type-1 *S*-loci [37]. Under the selective pressure from outcrossing, type-1 SI has been maintained in Plantaginaceae, Solanaceae, Rosaceae and Rutaceae by deletion or inactivation of the duplicate *S*-loci resulting from whole genome duplication (WGD) [37]. But in Brassicaceae, Papaveraceae, Primulaceae, Turneraceae, Linaceae and Oleaceae, according to our comprehensive analyses comprising all the recently reported SI systems, type-1 SI was lost either through *S*-locus deletions or by duplication maintenance, thus leading to the regain of the new family-specific SI system (type-2-5, 7 and 8) (Figure 1E). Since the ancient type-1 *S*-locus encoding Class I/II T2 RNase and FBA/FBK proteins originated from the MRCA of the angiosperms, it is possible that monocots have acquired type-6 SI following the loss of the ancient type-1 (Figure 1E; Supplemental Figure S2). In addition, we have summarized the molecular mechanisms of the eight SI types via cartoon diagrams (Figure 1F; Supplemental Figures S3-S9), with two reported working models of the widespread type-1 SI found in *Petunia hybrida* presented (Figure 1G) [38, 39].

### *S* gene identification and curation

The *S*-locus information of the assembled SI species has been fully investigated in PSIA (Figure 1H) based on the known molecular types. We started by downloading all known *S* genes from the NCBI Nucleotide and Protein Database and totally collected 1,275 nucleotide and 1,130 protein accessions of *S* genes from public resources. Next, we used blast+ to find *S* genes in all of the collected genome assemblies of SI plants [40], and manually curated the coding regions of each gene to ensure accurate splice sites. For *S* determinants containing conserved secondary domains, such as the F-box and FBA/FBK of SLF/SFB/SFBB and the Ribonuclease T2 family domain of S-RNase, we further searched for them to identify the candidate type-1 *S* [41]. For type-2 to type-8 SI, *S* genes were identified according to the blast results and linkage distance as previously described [33]. In total, 3,095 *S* genes have been identified and curated manually in PSIA.

### Useful Tools

User-friendly tools have been integrated into PSIA, including BLAST (genome blast), Sequenceserver 2.0 (*S* gene blast), Jbrowse, Synteny Viewer, and tools for phylogenetic analysis. ViroBlast has been incorporated into PSIA [42], and local blast databases have been established for several genome assemblies representing different SI types. Additionally, SequenceServer 2.0 has been implemented within PSIA and local blast databases have been developed using thousands of *S* genes [43], which is expected to significantly enhance the efficiency of SI research (Supplemental Figure S10). For genome assemblies with accessible GFF files, a Jbrowse function [44] has been provided, enabling researchers to browse the entire genome. The SynVisio (https://github.com/kiranbandi/synvisio) component will facilitate comparative genomic studies of different genome assemblies and *S*-loci. The synteny analyses of the *S*-loci will contribute to a deeper understanding of the dynamic evolution of SI (Supplemental Figure S11). Besides, phylogenetic trees have been constructed using IQtree based on the *S* genes identified in this study [45], which will aid in investigating their origins and relationships (Supplemental Figures S12 and S13).

## Perspectives and concluding remarks

The study of angiosperm self-incompatibility (SI) dates back to Darwin’s era, and while substantial progress has been made, much remains to be discovered about this fascinating reproductive mechanism. The growing availability of high-quality genomic sequences is a boon to systematic evolutionary analyses of *S* genes, offering deeper insights into the dynamic evolution of *S*-loci. PSIA stands as the first comprehensive knowledgebase dedicated to plant SI, consolidating all available genome assemblies of SI species from various sources, along with detailed *S*-locus information derived from these assemblies.

PSIA acts as a repository for SI-related information, including the origins and evolution of *S*-loci, the molecular mechanisms of SI, and representative SI models. Unlike other plant genome databases that often neglect *S* genes, PSIA comprehensively examines the *S*-locus information of assembled SI species. For example, while the Sol Genomics Network focuses on gene expression, quantitative trait loci (QTLs), and genetic markers within the Solanaceae family, PSIA enables efficient browsing, BLAST searching, analysis, and downloading of *S* genes across species exhibiting one of the eight known SI types.

PSIA also offers a comprehensive platform for comparative genomic studies of *S*-loci across different taxonomic levels, from genera to families and higher taxa, thus facilitating the exploration of the dynamic processes behind the origin, loss, and regain of SI.

In summary, PSIA is an essential knowledgebase for plant SI, making full use of *S*-locus sequences derived from rapidly advancing genome assembly technologies. Looking ahead, PSIA will be regularly updated to include the latest sequenced genomes and molecular mechanisms of plant SI. It is evident that PSIA will be an invaluable resource for exploring the molecular mechanisms of plant mating systems and for providing new insights into the evolutionary dynamics of *S*-loci.

## Supporting information

PSIA_Supplemental_Figures.pdf

S-genes-identified

## Authors’ contributions

YX and HZ conceived and supervised the project. CW collected and analyzed the SI data of Solanaceae, Rosaceae, Rutaceae, Plantaginaceae, Papaveraceae, Primulaceae, Turneraceae, Linaceae and Oleaceae. HKZ collected and analyzed the SI data of Poaceae. SS collected and analyzed the SI data of Brassicaceae. CW wrote the source code and constructed the database. HZ designed the molecular framework and the schematic diagram of the eight SI types. CW, HZ, and YX wrote the manuscript. All authors read and approved the final manuscript.

## Acknowledgments

This work was supported by the National Natural Science Foundation of China (32030007 and 32200273).

## Availability of data and materials

The datasets collected and analyzed during the current study are available on the website, [http://plantsi.cn/index.html].

