## Supplementary material for "PSI Atlas: a comprehensive knowledgebase of plant self-incompatibility": PSIA_Supplemental_Figures.pdf

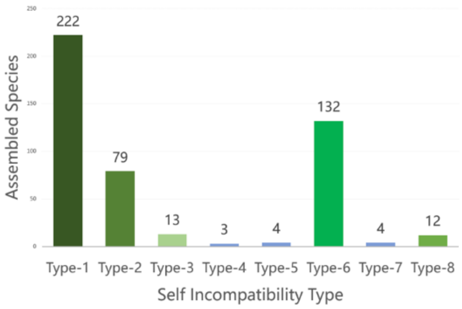


| Type | Count |
|---|---|
| SI Type | 8 |
| Family | 11 |
| Species | 469 |
| Genome Assemblies | 500+ |
| *S* Genes | 4370 |

**Supplemental Figure S1. General statistics of PSIA**

The number of assembled species related to eight types of SI has been summarized. The database will be kept updated along with the newly sequenced genomes of SI plants.

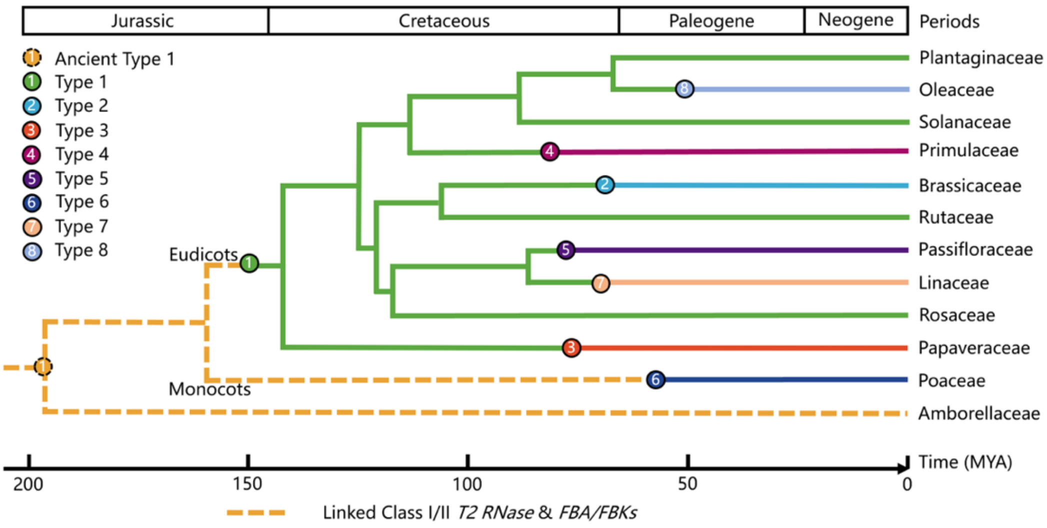


**Supplemental Figure S2. The phylogenetic tree of species at the family level represents self-incompatibility of 1-8 types.**

The evolutionary tree is generated by TimeTree. The geological time scale is at the top, while the bottom numerical axis represents the evolutionary timeline. The circles of different colors and their corresponding lines represent the eight types of SI. The orange dashed line indicates the ancient type 1 S structure.

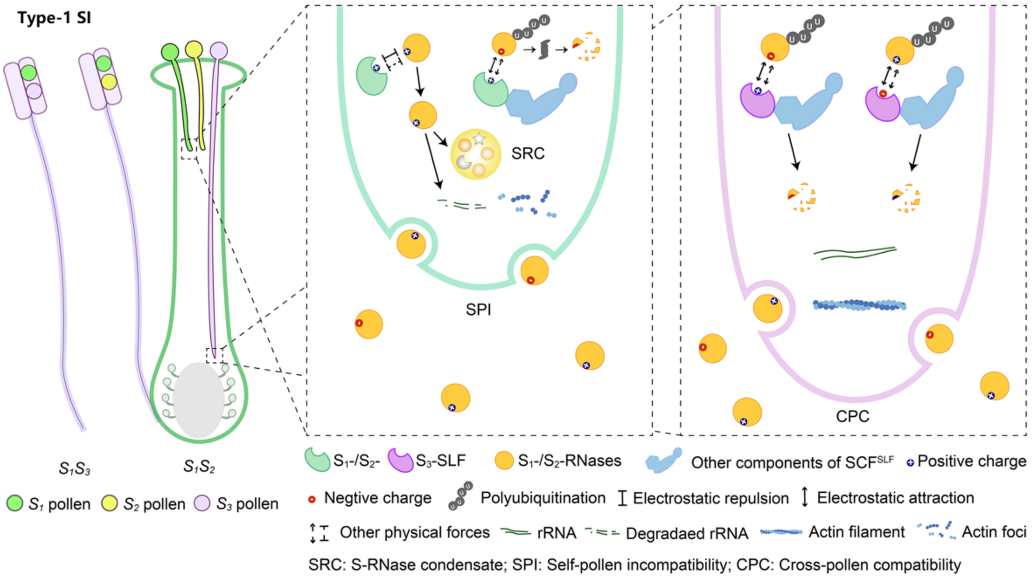


**Supplemental Figure S3. Molecular mechanism of type-1 SI.**

The system with the broadest taxonomic distribution, which we term type-1 SI, is gametophytic and based on linked pistil and pollen *S* components, corresponding to *S-RNase* and *S-locus F-box* (*SLF*), also named *S-haplotype-specific F-box* (*SFB*), respectively. So far, type-1 SI has been found in four eudicot families: Solanaceae, Plantaginaceae, Rosaceae, and Rutaceae, spanning two major clades (superrosids and superasterids).

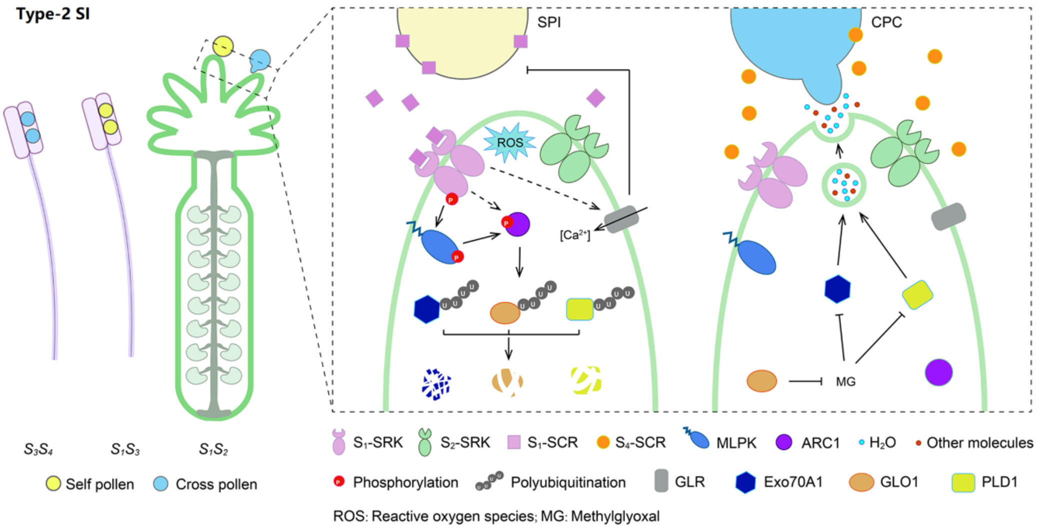

**Supplemental Figure S4. Molecular mechanism of type-2 SI.**

Type-2 SI is the sporophytic Brassicaceae-type SI, controlled by a male *S*-locus cysteine-rich (SCR) protein/*S*-locus protein 11 and a female *S*-locus receptor kinase (SRK).

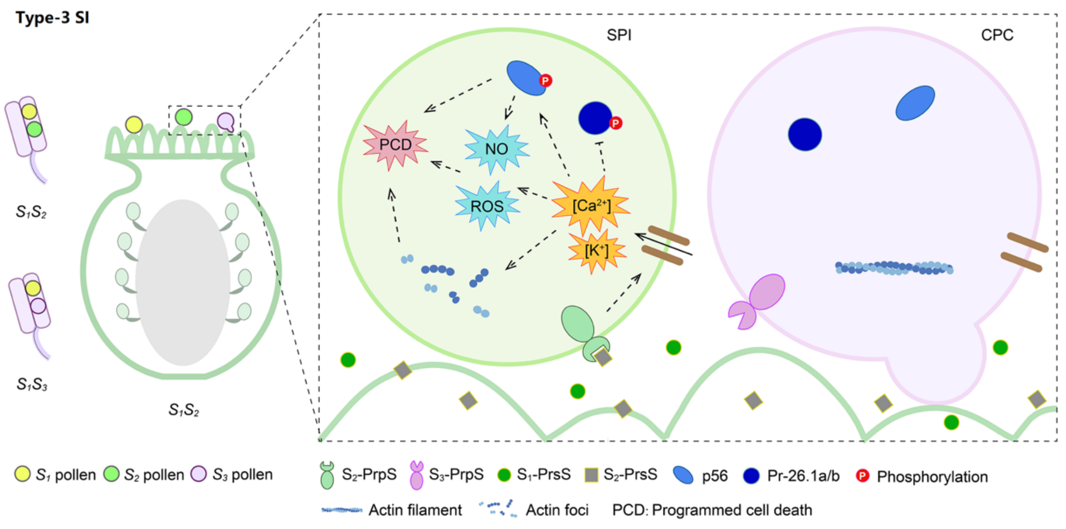


**Supplemental Figure S5. Molecular mechanism of type-3 SI.**

Type-3 is the gametophytic Papaveraceae-type SI, possessing the common poppy (*Papaver rhoeas*) stigma *S* (PrsS) and *P. rhoeas* pollen *S* (PrpS).

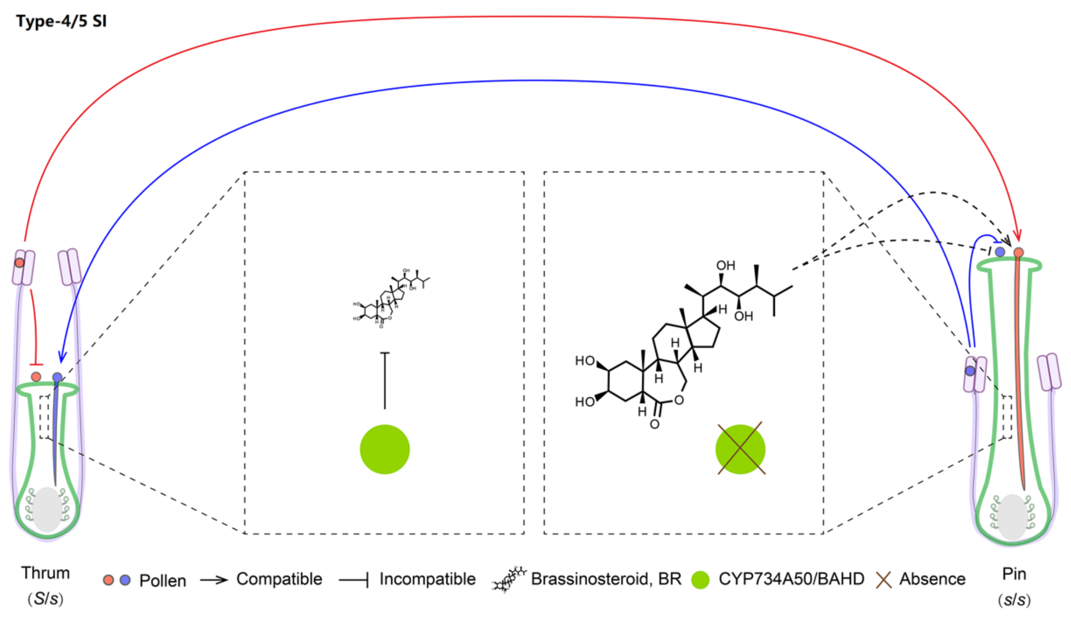


**Supplemental Figure S6. Molecular mechanism of type-4/5 SI.**

Type-4 is the sporophytic heterostyly of *Primula*, involving the *S*-locus supergene consisting of five genes encoding style length-determining cytochrome P450 (CYP450), anther position-controlling GLOBOSA (GLO), a functionally unknown Conserved Cysteine Motif (CCM), Pumilio-like RNA-binding protein (PUM) and a Kelch repeat F-Box (KFB).

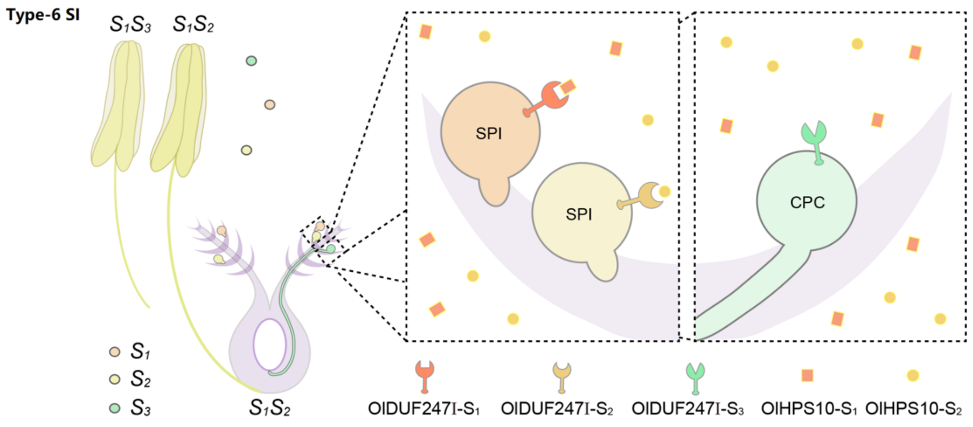


**Supplemental Figure S7. Molecular mechanism of type-6 SI.**

We classified the self-incompatibility of poaceae as type-6. In the grass family (Poaceae), which contains all the cereal and major forage crops, SI has been known for half a century to be controlled gametophytically by two multiallelic and independent loci, *S* and *Z*. Evidence showed that both the *S* and *Z* genes encode two stamen-expressed polypeptides containing a domain of unknown function 247 (DUF247) (*S*- and *Z-DUF247s*) and a pistil-specific one possessing a signal peptide (*sS/sZ*).

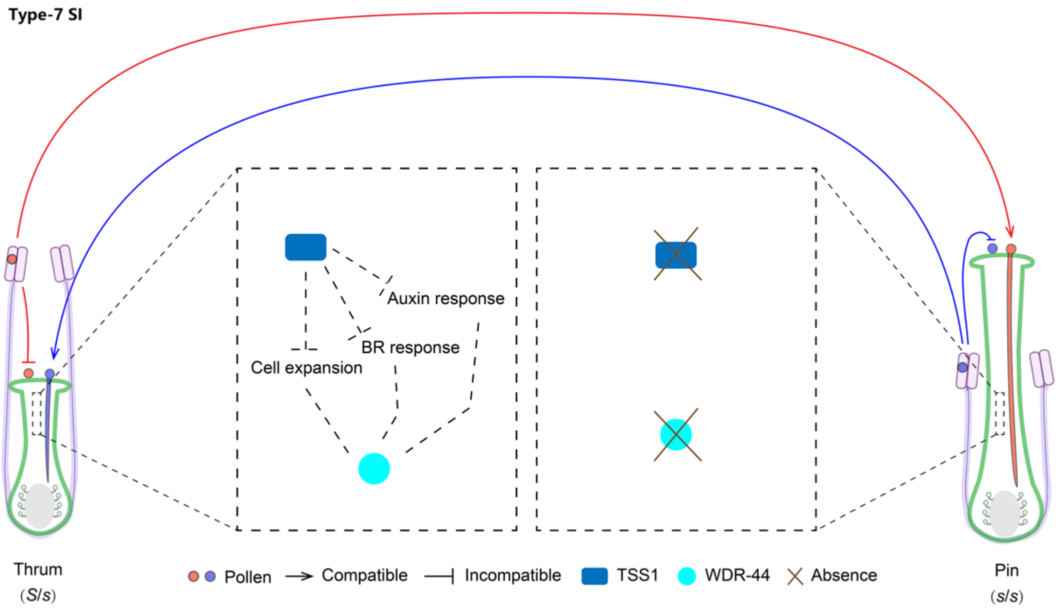


**Supplemental Figure S8. Molecular mechanism of type-7 SI.**

We classified the self-incompatibility of Linaceae as type-7. The *S*-locus of *Linum tenue* was identified using population genomic data. The results show that hemizygosity and thrum-specific expression of *S*-linked genes, including a pistil-expressed candidate gene for style length, are major features of the Linum *S*-locus. Two distyly candidate genes (*LtTSS1* and *LtWDR-44*) are only present in the dominant *S* allele.

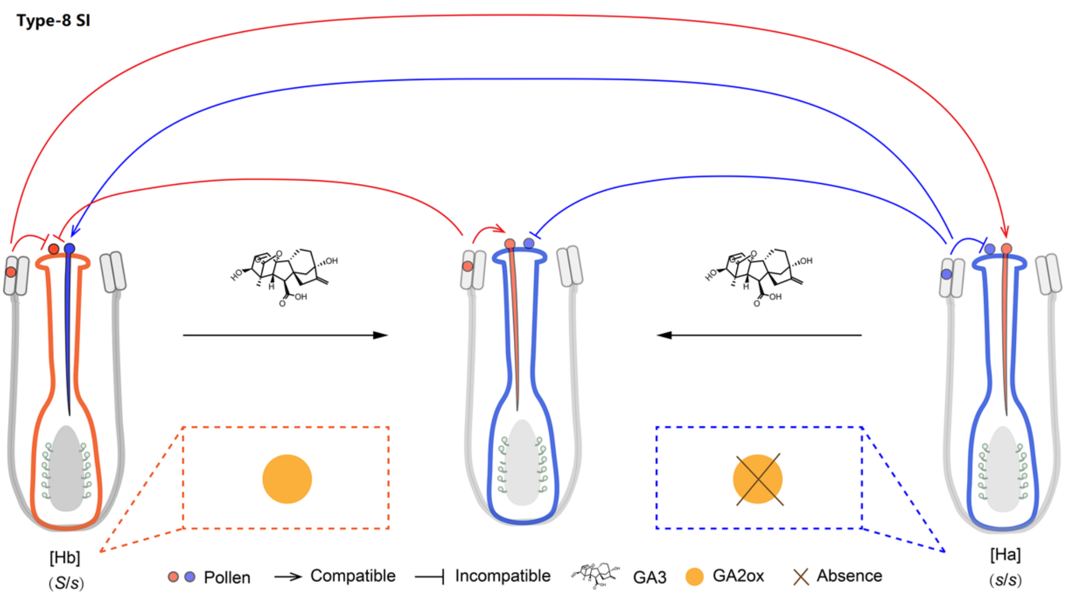


**Supplemental Figure S9. Molecular mechanism of type-8 SI.**

We classified the self-incompatibility of Oleaceae as type-8. The *S*-locus region of *Phillyrea angustifolia* proved to have a segregating 543-kb indel unique to one specificity, suggesting a hemizygous region, as observed in all distylous systems so far studied at the genomic level. Only one of the predicted genes (*GA2ox-S*) in this indel region is found in the olive tree. The presence/absence polymorphism of the *GA2ox-S* gene is stably associated with SI groups across Oleaceae.

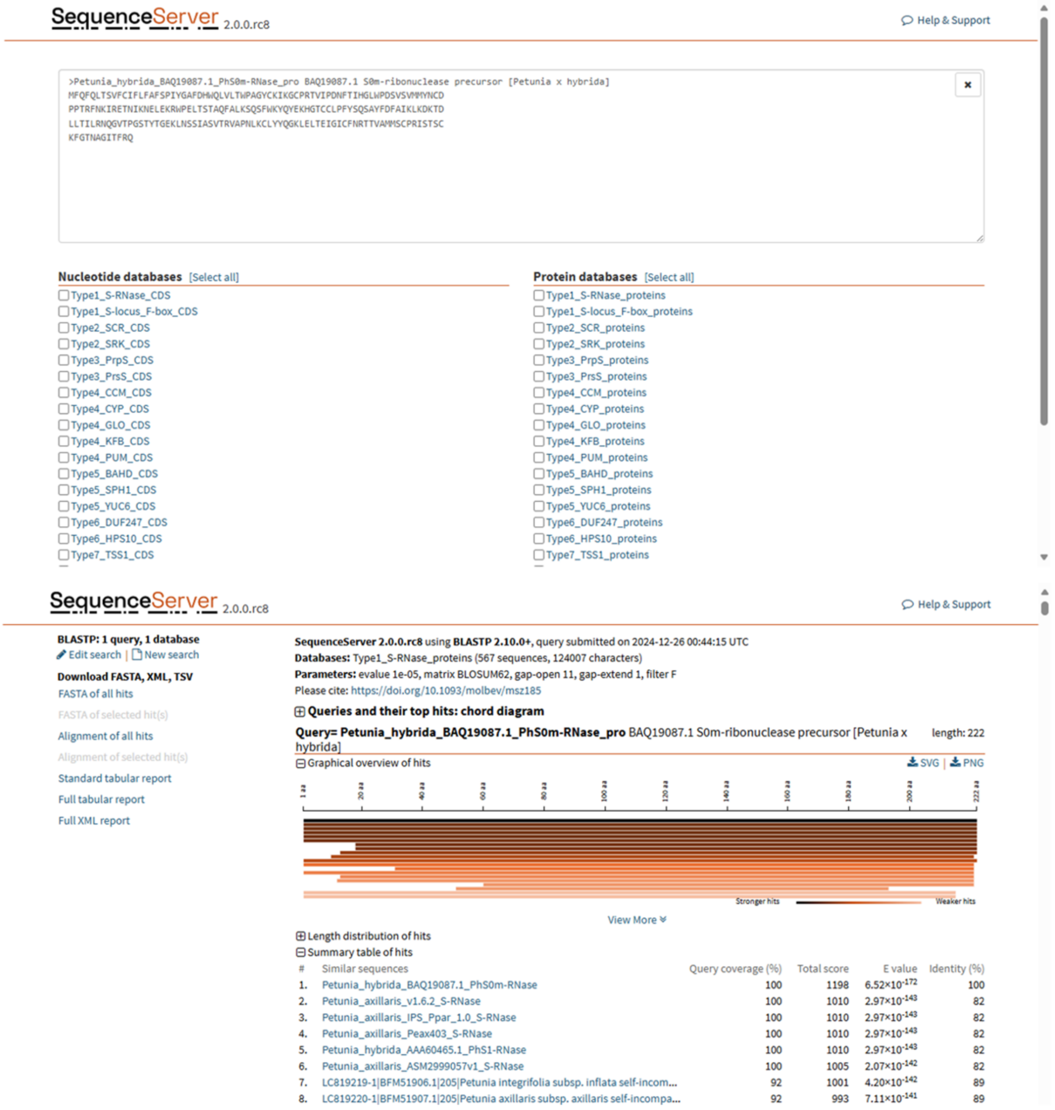


**Supplemental Figure S10. BLAST against almost the most complete repertoire of S genes of different SI types.**

The SequenceServer was integrated into PSIA, and we constructed local blast databases using thousands of S genes (publicly known and newly identified S genes), which will greatly improve the efficiency of SI research.

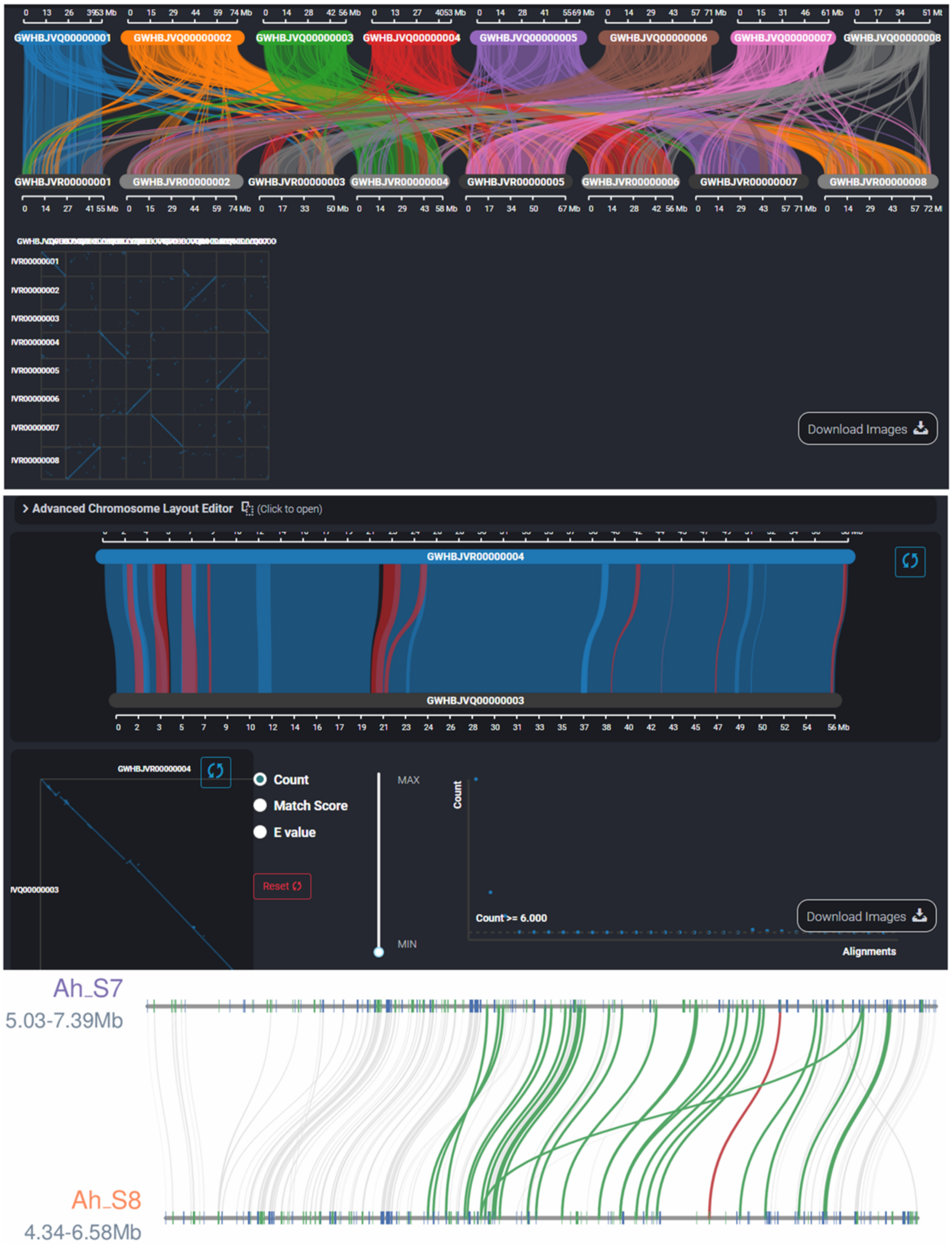


**Supplemental Figure S11. Synteny Viewer tool of PSIA.**

Top, the results of comparative genomics between different genome assemblies. Middle, the synteny between the chromosomes of *S*-locus. Bottom, the synteny between the *S*-locus.

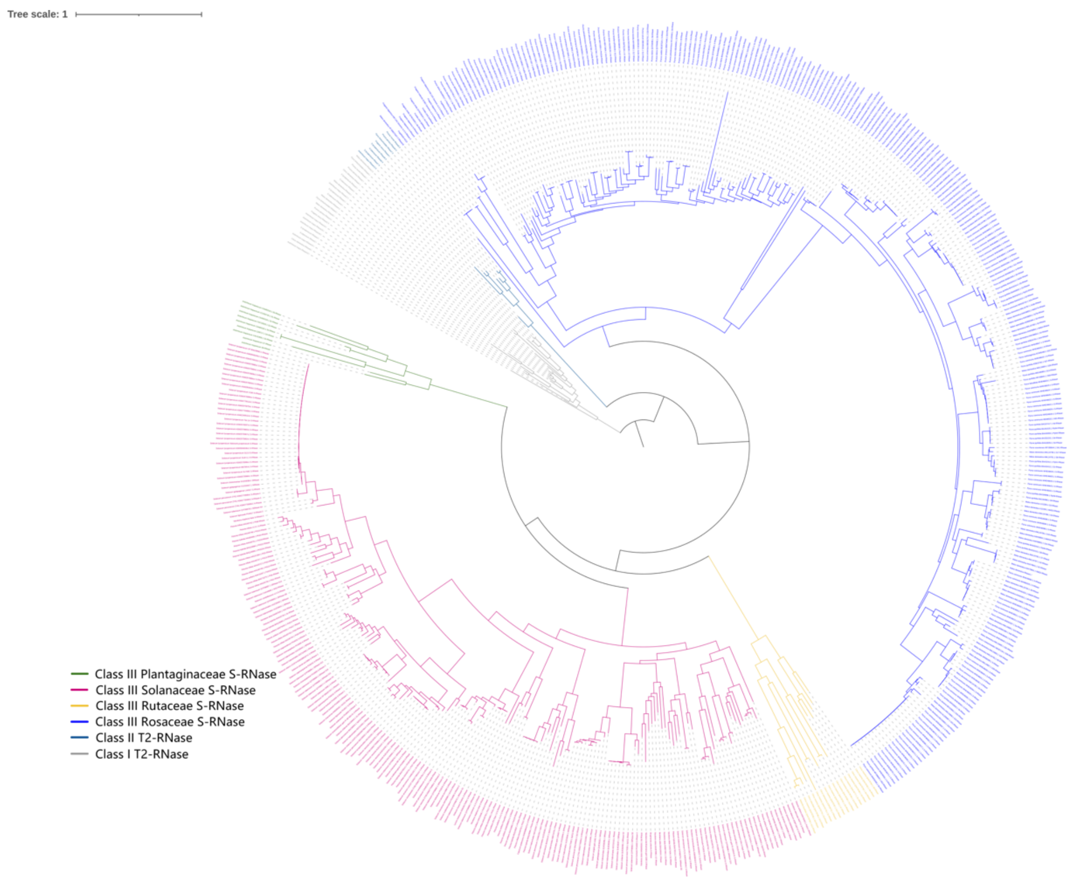


**Supplemental Figure S12. Maximum-likelihood tree of the S-RNases of the four type-1 SI families.**

Maximum-likelihood tree of the S-RNases. S-RNases (Class III T2 RNases) from four families (Plantaginaceae, Solanaceae, Rutaceae, Rosaceae) and other types of T2 RNases (Class I and II) are indicated by different branch colors.

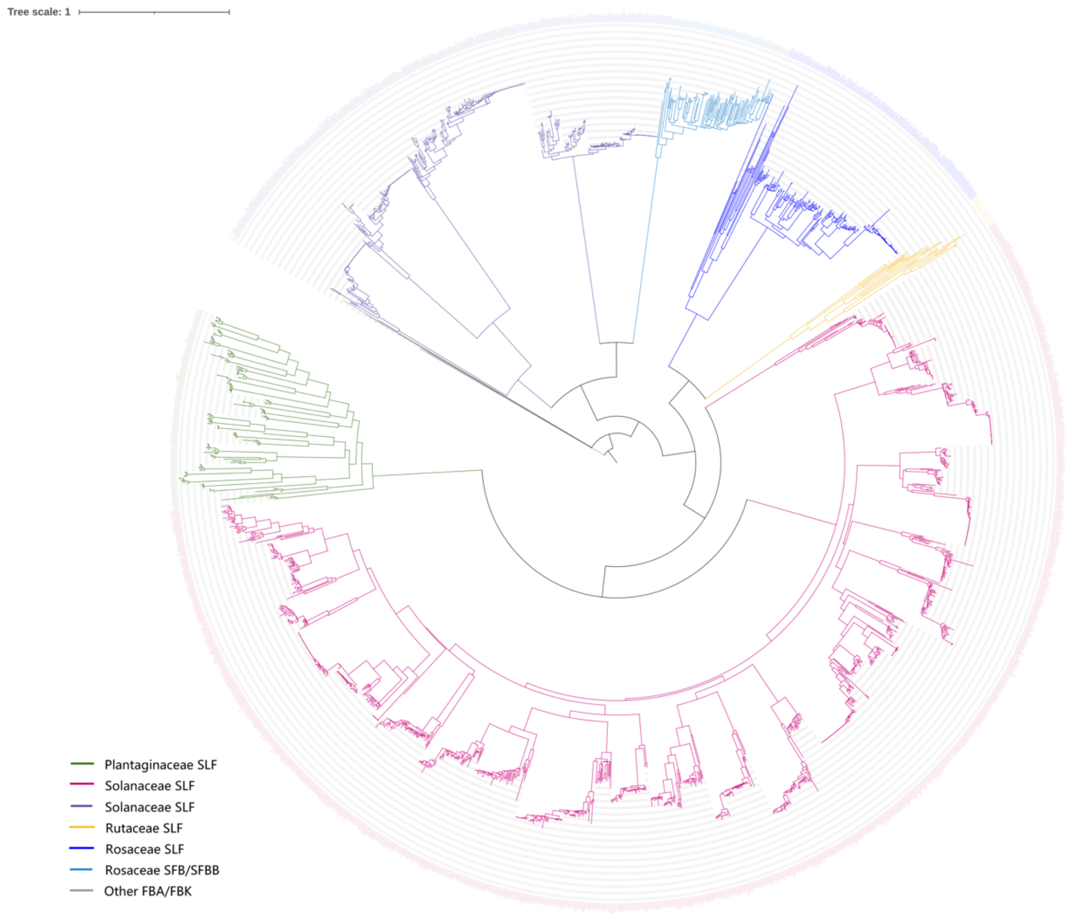


**Supplemental Figure S13. Maximum-likelihood tree of the SLFs/SFBs/SFBBs of four type-1 SI families.**

Maximum-likelihood tree of the SLFs/SFBs/SFBBs. SLFs/SFBs/SFBBs from four families (Plantaginaceae, Solanaceae, Rutaceae, Rosaceae) and other FBA/FBKs are indicated by different branch colors.
